# Retromer-mediated recruitment of the WASH complex involves discrete interactions between VPS35, VPS29 and Fam21

**DOI:** 10.1101/2024.01.22.576619

**Authors:** Miguel Romano-Moreno, Elsa N Astorga-Simón, Adriana L Rojas, Aitor Hierro

## Abstract

Endosomal trafficking ensures the proper distribution of lipids and proteins to various cellular compartments, facilitating intracellular communication, nutrient transport, waste disposal, and the maintenance of cell structure. Retromer, a peripheral membrane protein complex, plays an important role in this process by recruiting the associated actin-polymerizing WASH complex to establish distinct sorting domains. The WASH complex is recruited through the interaction of the VPS35 subunit of retromer with the WASH complex subunit FAM21. Here, we report the identification of two separate fragments of Fam21 that interact with VPS35, along with a third fragment that binds to the VPS29 subunit of retromer. The crystal structure of VPS29 bound to a peptide derived from Fam21 shows a distinctive sharp bend that inserts into a conserved hydrophobic pocket with a binding mode similar to that adopted by other VPS29 effectors. Interestingly, despite the network of interactions between Fam21 and retromer occurring near the Parkinson’s disease-linked mutation (D620N) in VPS35, this mutation does not significantly impair the direct association with Fam21 *in vitro*.

## INTRODUCTION

Retromer is a protein complex that promotes the selective retrieval and recycling of hundreds of integral membrane proteins, referred to as cargo, from endosomes back to the *trans*-Golgi network, the cell surface, or other specialized organelles. The impairment of retromer function has indeed been associated with numerous pathologies, particularly neurodegenerative disorders, including Alzheimer’s disease (AD) (48; 36), Parkinson’s disease (PD) (55), Down’s syndrome (DS) (7), and amyotrophic lateral sclerosis (ALS) (39; 41). Not surprisingly, there is a growing interest in exploring retromer as a target for therapeutic interventions (35; 39; 3). The mammalian retromer consists of an evolutionary conserved trimer of vacuolar protein sorting (VPS) 35, VPS29, and VPS26, which orchestrates cargo selection, membrane deformation, and cytoskeletal association through interactions with accessory proteins. The function of the retromer is linked with its recruitment to the endosomal membrane, mediated by a subset sorting nexin (SNX) proteins (reviewed in (50; 53), along with Rab5 and Rab7 GTPases (27). Once recruited to endosomes, retromer facilitates selective cargo sorting into distinct transport carriers by forming actin-decorated microdomains (43; 49).

Actin polymerization contributes to many membrane remodeling processes including endocytosis, endosomal recycling, autophagy and vesicle trafficking. An important component in vesicle dynamics is the Arp2/3 complex which controls actin nucleation and branching of filaments for the generation of membrane subdomains, as well as mechanical forces during vesicle formation and scission. Proper functioning of the Arp2/3 complex requires members of the Wiskott-Aldrich Syndrome Protein (WASP) family, termed Nucleation-Promoting Factors (NPFs), for the spatial and temporal organization of actin filaments. The WASP and SCAR Homolog WASH is the major NPF at the surface of endosomes and is an integral part of a pentameric complex which includes WASH (WASHC1), the family with sequence similarity 21 (FAM21 or WASHC2), the coiled coil domain containing protein 53 (CCDC53 or WASHC3), Strumpellin (WASHC5), and the Strumpellin and WASH interacting protein (SWIP or WASHC4) (21). WASH complex exhibits significant homology to another WASP family member, the WAVE regulatory complex (WRC) found at the plasma membrane (21). Nonetheless, the overall resemblance between WRC and the WASH complex is only applicable to four of their subunits (21). The major difference corresponds to FAM21 which is a much larger protein (∼1300 amino acids) compared to its counterpart subunit (ABI, ∼370 amino acids) in the WRC complex. FAM21 is formed by a small (∼200 amino acids) ‘head’ domain required for interaction with other WASH complex subunits and a long (∼1100 amino acids) ‘tail’, mostly unstructured, that mediates direct biding to several proteins including the acting-capping proteins CAPZa and CAPZb (17), the DNAJC13 protein, known as receptor-mediated endocytosis-8 (RME-8) (12), the CCDC22 and CCDC93 subunits of the CCC (COMMD/CCDC22/CCDC93) complex (14; 42), the FK506-binding protein 15 (FKBP15) (14; 40), the cargo adaptor SNX27 (49; 29), and the VPS35 subunit of the retromer complex (14; 20; 16) (Figure 1A). This network of interactions suggests different mechanisms for controlling actin polymerization on endosomes (47). Yet, the localization of the WASH complex to endosomes largely relies on the interaction between FAM21 and retromer (13; 14; 16). The large unstructured C-terminal tail of FAM21 contains 21 repeats of the consensus L- F-[D/E]_3-10_-L-F sequence, leucine-phenylalanine-acidic (LFa) motif, that bind via multivalent interactions to VPS35 (20). Nonetheless, despite the general similarity among the 21 repeats, only those at the very C-terminal end of the tail significantly contribute to WASH complex recruitment (20; 16). In addition, the interaction between FAM21 and VPS35 requires the presence of VPS29, suggesting a more complex pattern of interactions than initially anticipated (16; 45). Interestingly, genetic studies of sporadic and familial forms of Parkinson’s disease (PD) identified a point mutation in VPS35 (D620N) as the causative factor in the development of the disease (51; 55). The D620N mutation does not affect the stability or assembly of VPS35 with other retromer subunits but impairs its association with FAM21 and the recruitment of the WASH complex to endosomes (34; 54; 6). The ankyrin-repeat-domain-containing protein 50 (ANKRD50) is another retromer-SNX27-SHRC interacting protein that fails to associate with VPS35 harboring the D620N mutation (26). These alterations lead to altered endosomal morphology (6) and impaired sorting of some retromer cargos such as the cation-independent mannose 6-phosphate receptor (CI-MPR), the post-synaptic AMPA receptor GluR1, and the Autophagy-related protein 9A (ATG9a), but not retromer-SNX27 cargo like the Glucose transporter 1 (GLUT1) (11; 34; 54; 37; 10; 6). Intriguingly, while VPS35 interacts robustly with endogenous Fam21 in cell culture models, the equivalent interaction has not been observed using similar co-immunoprecipitation (co-IP) assays from mouse brain extracts suggesting that it may form a distinct tissue-specific network of interactions or a labile complex (4). While these findings suggest a delicate interplay between retromer and the WASH complex to control actin dynamics, most molecular details regarding how retromer interacts with FAM21 remain unknown.

**FIGURE 1:**
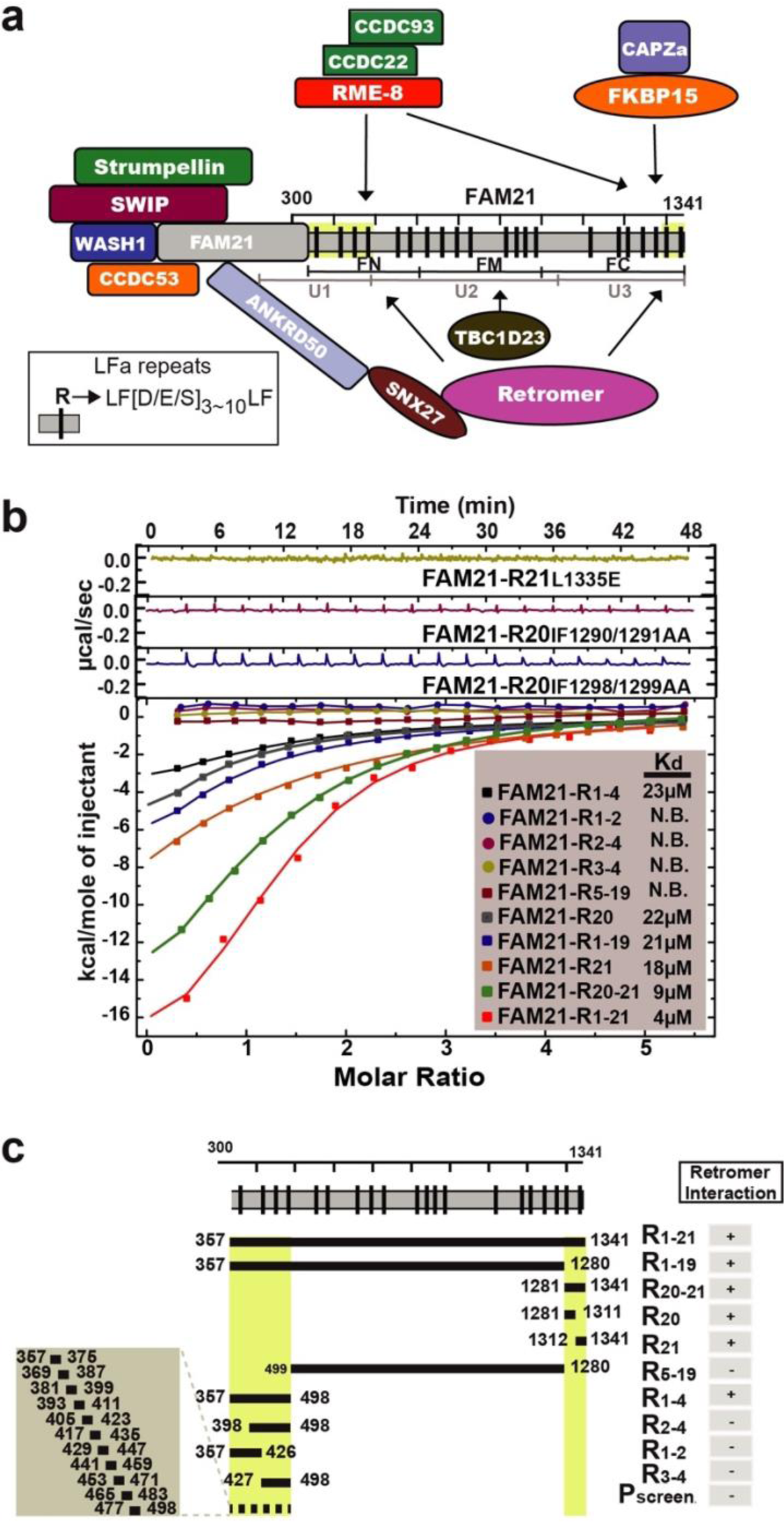
The long, disordered tail of Fam21 interacts with retromer through three regions. (a) Schematic cartoon depicting the association between various endosomal proteins and the Fam21 tail. Vertical black bars within the extended tail represent each of the 21 LFa repeats. (b) Representative ITC measurements illustrate the interaction of distinct segments of the Fam21 tail titrated into retromer. The top three panels correspond to the titration curves for the indicated mutants within the R_20_ or R_21_ LFa motifs. The lower panel illustrates the binding isotherms for each fragment of Fam21. Raw titration curves for the binding isotherms and the thermodynamic parameters are summarized in Figure S1 and Table S1, respectively. (c) Schematic representation of the Fam21 constructs used, with green areas indicating the binding regions towards retromer.

In the present study, we provide a detailed biochemical and structural characterization of the interactions between retromer and FAM21, required for the endosomal association of the WASH complex. We demonstrate that FAM21 harbors two distinct binding sites toward VPS35 and a third one toward VPS29 within a conserved pocket shared by other cellular ligands. The network of FAM21 interactions occurs in proximity to the apex of the arch formed by retromer dimers. Interestingly, the Parkinson’s VPS35(D620N) mutation, situated near the VPS35:VPS35 interface, does not significantly reduce the association with Fam21 *in vitro*.

## RESULTS

### Retromer complex interacts with distinct segments of the Fam21 tail

The 21 repeats (R1 to R21) of the LFa motif are distributed along the long unstructured tail of FAM21(20). However, the division of the tail into different fragments have shown strong variations with respect to their contributions to retromer binding (20; 16). These findings suggest a more complex recognition mechanism than initially anticipated, possibly directed by multiple interactions and binding modes. To define more precisely the interactions between FAM21 and retromer we first confirmed several of the previously described interactions by isothermal titration calorimetry (ITC), and then expanded the analysis to new engineered fragments encompassing distinct LFa motifs. We generated five main fragments harboring multiple LFa motifs. The largest fragment corresponded to the entire tail region with the 21 repeats of the LFa motif (denoted as R_1- 21_) whereas shorter fragments included R_1-19_, R_20-21_, R_1-4_ and R_5-19_ (Figure 1B, C). The ITC analysis confirmed direct interactions of two wide apart regions of FAM21, R_20-21_ and R_1-4_ with retromer (Figure 1B, C, and Supplementary Figure 1A). In contrast, the large middle region R_5-19_ did not yield any measurable interaction in solution. Out of the two regions that interact with retromer, the R_20-21_ exhibited the strongest contribution to the binding in agreement with previous studies (20; 16). To further delineate these interactions, we narrowed the search using synthetic and recombinantly expressed peptides covering R_1-4_ and R_20-21_ regions (Figure 1B, C). Interestingly, while R_20_ and R_21_ fragments exhibited significant binding to retromer, neither of the peptides encompassing the R_1-4_ region, nor the fragments R_1-2_, R_2-4_ or R_3-4_ showed any interaction (Figure 1B, C and S1 ). Indeed, the interactions of R_20_ and R_21_ with retromer were abolished by single or double alanine substitutions within their respective LFa motifs which evidenced for a direct physical interaction by short sequence patches (Figure 1B, C). These observations hinted at distinct binding modes between the two stretches. In this sense, whilst R_20_ and R_21_ functioned as independent linear motifs, the R_1-4_ region seemed to rely on conformational constraints for binding.

### Fam21 interacts with retromer through VPS35 and VPS29 subunits

We next aimed at identifying the retromer sites to which R_1-4_, R_20_ and R_21_ bind. First, considering that retromer is an elongated complex where the VPS26 and VPS29 subunits associate with the N-terminal and C-terminal ends of the VPS35 subunit, respectively, we divided VPS35 into roughly two halves: VPS35N (aa 14-470) and VPS35C (aa 476-780). This division allowed us to produce two subcomplexes, namely the VPS26–VPS35N subcomplex and the VPS29–VPS35C subcomplex. We then evaluated their affinities towards R_1-4_, R_20_, and R_21_. We found that R_1-4_ and R_21_ bound to VPS29–VPS35C subcomplex with similar affinity as to the full retromer, but unexpectedly, R_20_ did not interact with any of the subcomplexes (Figure 2A-D, E and S2). We reasoned that R_20_ could bind the middle section of VPS35 that was not included in the constructs. To test this possibility, we designed an extended VPS35C construct (aa 204-780) and, as expected, the R_20_ bound to the VPS29–VPS35_204-780_ subcomplex with equivalent affinity as to the whole retromer. Next, we analyzed the interaction of R_1-4_ and R_21_ with each component of the VPS29–VPS35C subcomplex. Remarkably, R_1-4_ exhibited binding to VPS35C, whereas the R_21_ associated with VPS29, indicating distinct binding subunits for each region.

**FIGURE 2:**
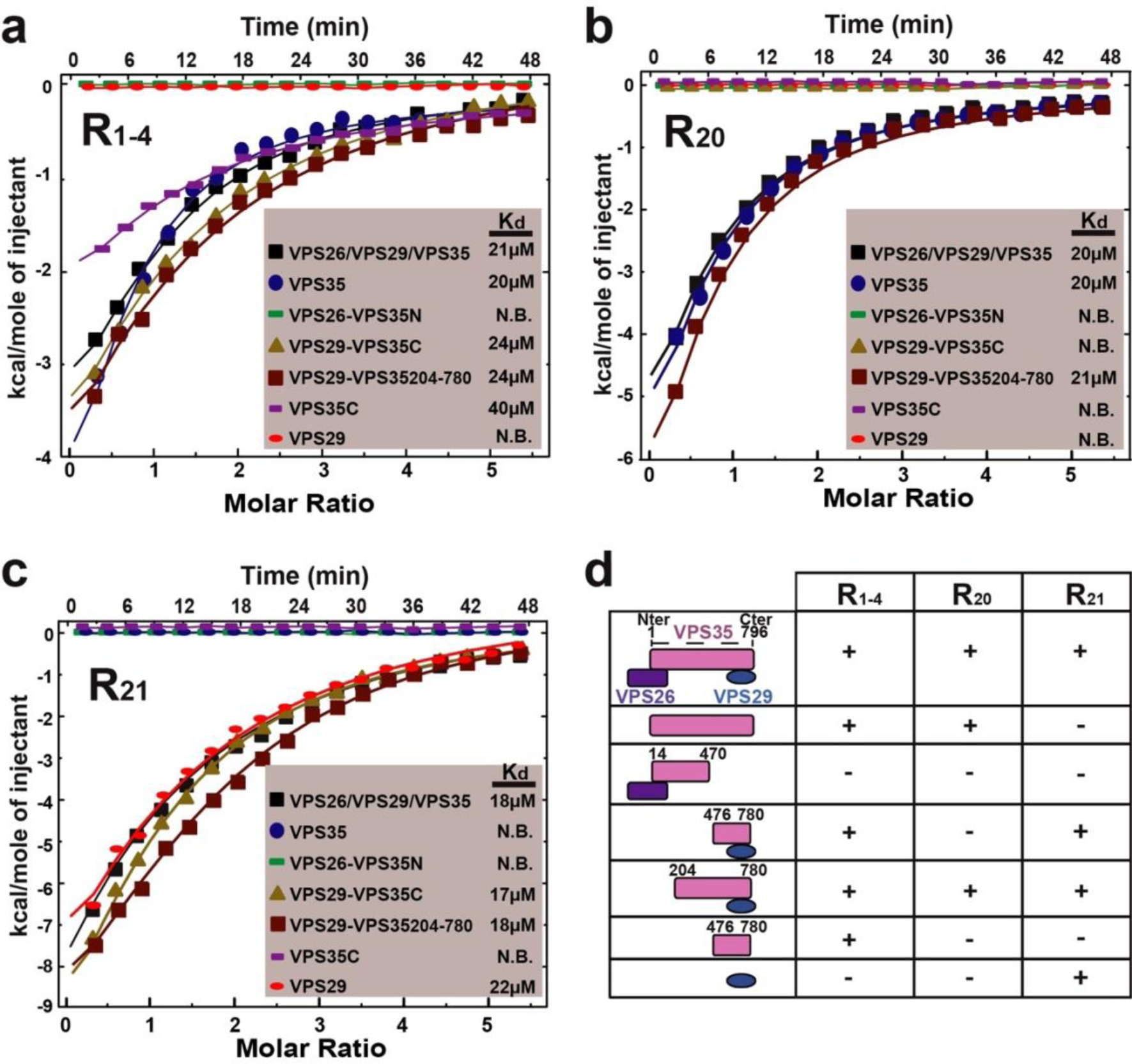
Mapping Fam21-interacting fragments on retromer. (a) Binding isotherms for the R_1-4_ fragment, (b) R_20_ fragment, and (c) R_21_ fragment to individual retromer subunits or truncated subcomplexes. Raw titration curves and thermodynamic parameters are summarized in Figure S1 and Table S1, respectively. (d) Schematic representation of the retromer subunits and truncated subcomplexes used for mapping the interactions.

### Identification of R_1-4_, R_20_ and R_21_ partner-binding surfaces on VPS35 and VPS29

Our previous experiments narrowed the interaction of the FAM21 tail down to two regions in VPS35 and one site on VPS29. Next, to identify amino acid residues important for the interaction, we selected several double and triple mutants on the basis of their solvent accessibility, conservation and spatial proximity in the three-dimensional structure. For mapping the R_1-4_-binding site in VPS35 we selected the mutants KL659/661AE, EP643/648AA, and KKK544/548/552AAA (Figure 3A). The only mutant that significantly altered the affinity for R_1-4_ was KL659/661AE (Figure 3A, B and S3). Similarly, for mapping the R_20_-binding site in VPS35 we evaluated the substitutions YI447/451AA, LK508/515AA, and PPV472/475/476AAA. In this case, only the substitution YI447/451AA precluded the interaction with R_20_ (Figure 3A, C). Finally, to map the region within VPS29 responsible for binding to R_21_, we focused on a conserved pocket located on the opposite side from the contact with VPS35. This pocket has been shown to be important for binding to the Rab7 GTPase activating protein TBC1 (Tre-2/USP6, BUB2, Cdc16) Domain Family, Member 5 (TBC1D5) (22), the VPS9- domain ankyrin repeated protein (VARP) (5), the Vacuolar Protein Sorting 35-Like (VPS35L) (15), and the *Legionella pneumophila* effector protein RidL (2; 44; 52). Thus, we introduced mutations to various residues within the conserved pocket of VPS29, including L152E, Y163A, and Y165A, known to play a crucial role in interactions with TBC1D5, VARP, VPS35L, and RidL. Notably, all three mutations proved indispensable for the interaction with R_21_ (Figure 2A, D), highlighting a shared binding pocket in VPS29 among multiple retrieval complexes.

**FIGURE 3:**
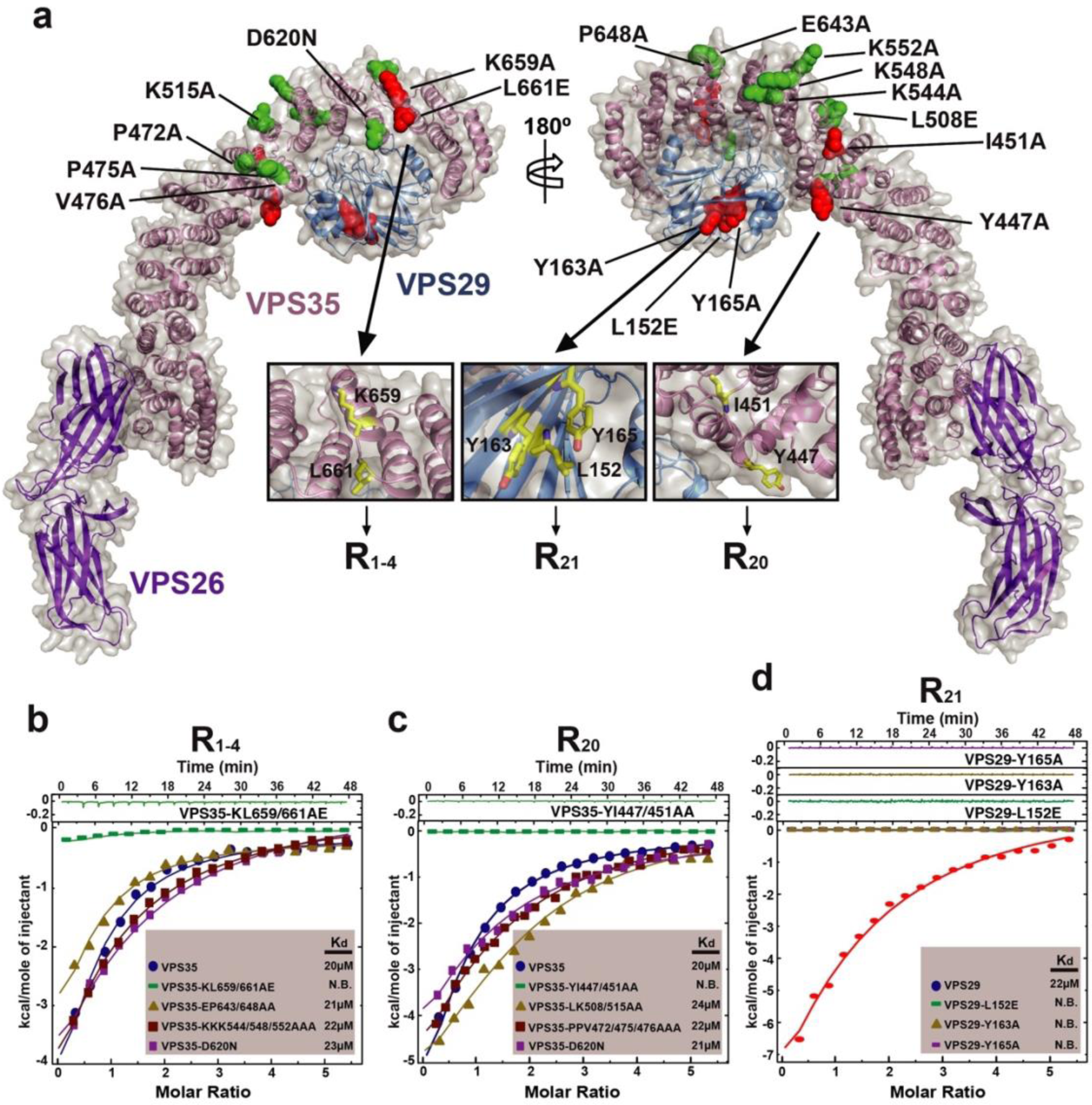
Mapping of Fam21’s binding sites on retromer using a structurally guided site-directed mutagenesis approach. (a) Semi-transparent surface overlaying a cartoon representation of full-length retromer (PDB 6VAC (24)). Mutated residues are shown in a ball-and-stick representation, color-coded in red for mutations interfering with binding and green for mutations not affecting binding towards the respective Fam21 fragments. (b-d) ITC analysis showing the binding of (b) R_1-4_ fragment, (c) R_20_ fragment, and (d) R_21_ fragment to both wildtype retromer subunits and selected mutants. Raw titration curves and thermodynamic parameters are summarized in Figure S1 and Table S1, respectively.

Previous studies have suggested that the VPS35(D620N) mutation linked to Parkinson’s disease impairs the recruitment of the pentameric WASH complex to endosomes (34; 54). This impairment was attributed to a reduced interaction between VPS29–VPS35C and the R_20-21_ motifs of FAM21 (34). Given the close proximity between D620 and the binding area for R_1-4_ (Figure 3A), we investigated whether D620N could influence the association with R_1-4_ using ITC. The results indicated that the retromer complex bearing the VPS35(D620N) mutation exhibited no significant differences compared to the wild type in binding to either R_1-4_ or R_20_ (Figure 3B, C). We did not assess the binding towards R_21_, as it binds to Vps29, which is far from the D620N site (Figure 3A).

### Structure of VPS29 bound to the R_21_ motif of FAM21

Previous crystallographic studies have solved the structure of retromer from two subcomplexes, the C-terminal region of VPS35 (VPS35C, amino acids 476–780) bound to VPS29 (19), and the N-terminal region of VPS35 (VPS35N, amino acids 14-470) bound to VPS26 and SNX3 (31). Taking advantage of the crystallization ability of these constructs we attempted to co-crystallize R_1-4_, R_20_ and R_21_ with their corresponding retromer subcomplex. Unfortunately, despite extensive efforts we were unable to obtain crystals of R_1-4_ and R_20_ in complex with retromer. On the contrary, we succeeded on the crystallization of R_21_ with VPS35C-VPS29. We determined the crystal structure by molecular replacement using the coordinates of VPS35C-VPS29 (PDB 2R17) (19) as the search model, and refined to 2.97 Å resolution with R_factor_ and R_free_ of 25.08 and 28.77, respectively (Table 1). The difference Fourier map clearly showed a portion of R_21_ comprising amino acids _1331_FDDPLNA_1337_ (Figure 4A). The remaining residues were not modeled due to missing electron density which most probably reflected intrinsic disorder. Overall, the central region of R_21_ forms a sharp turn inserted into the conserved pocket of VPS29 (Figure 4B and S4A, B). In agreement with the biochemical data described before, R_21_ anchors to VPS29 by the burial of P1334 and L1335 into a hydrophobic cavity formed by L2, L25, L26, F150, L152 and Y163. More peripherally F1331 of R_21_ stacks against the R176 aliphatic side-chain of VPS29 (Figure 4B, C). In addition, the interface includes a network of sidechain and backbone hydrogen bonds that further stabilize the association between R_21_ and VPS29 (Figure 4C). In this regard, the crystal structure provides key structural determinants for the interaction of R_21_ with VPS29.

**FIGURE 4:**
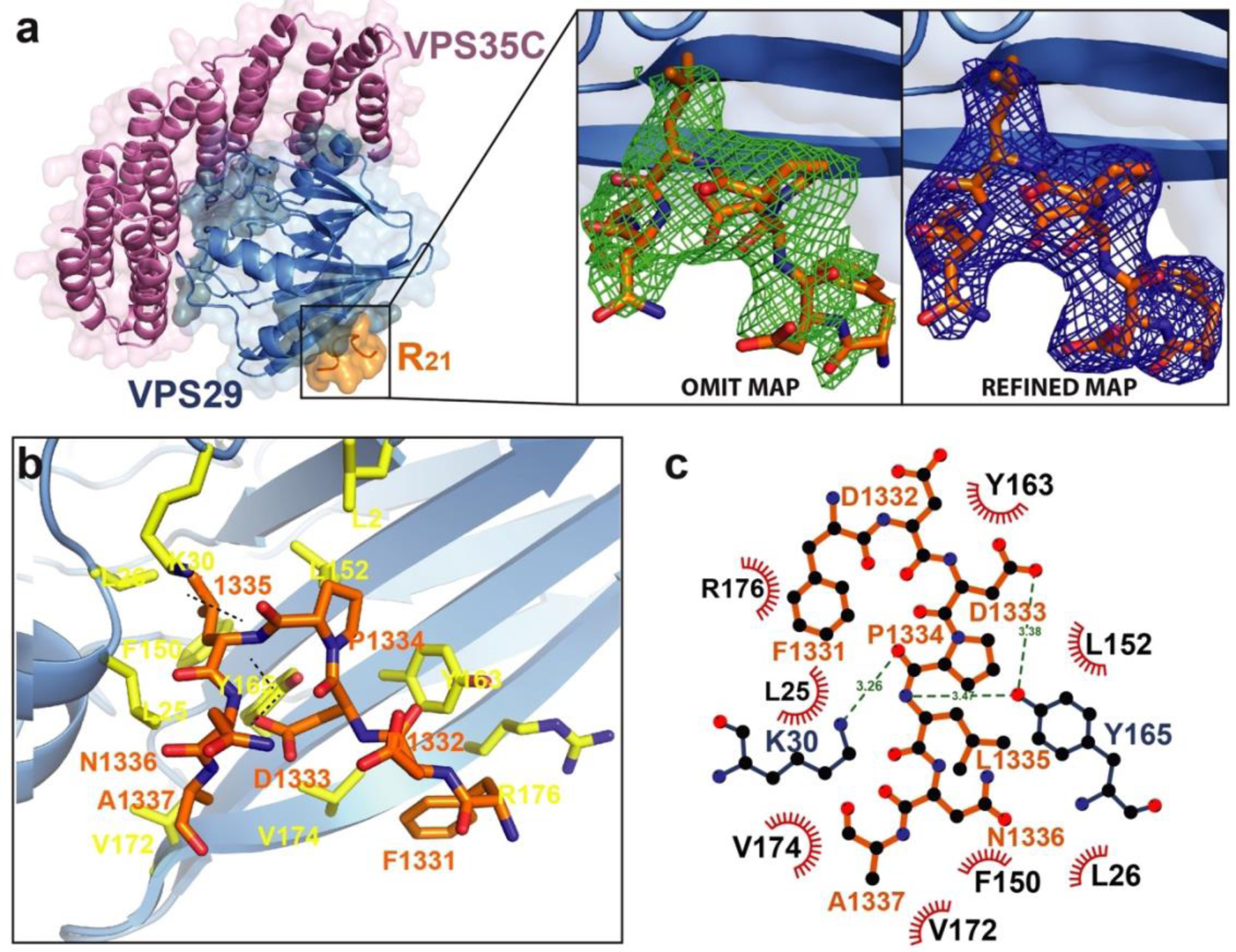
Crystal structure of R_21_ bound to retromer. (a) The overall structure of the complex formed by the R_21_ motif and VPS29-VPS35C in cartoon representation with transparent surfaces. Left insets show the omit difference electron density map (Fo-Fc) in green, contoured at 2.0σ, and the final refined 2Fo-Fc electron density map in blue, contoured at 1.5σ. (b) Detailed view of the R_21_-VPS29 interaction with relevant residues shown as sticks. (c) Interaction diagram generated with LigPlot+ (28), illustrating hydrophobic contacts, hydrogen bonds, and solvent accessibility of the R21 (aa 1331- FDDPLNA-1337) binding motif.

**TABLE 1:**
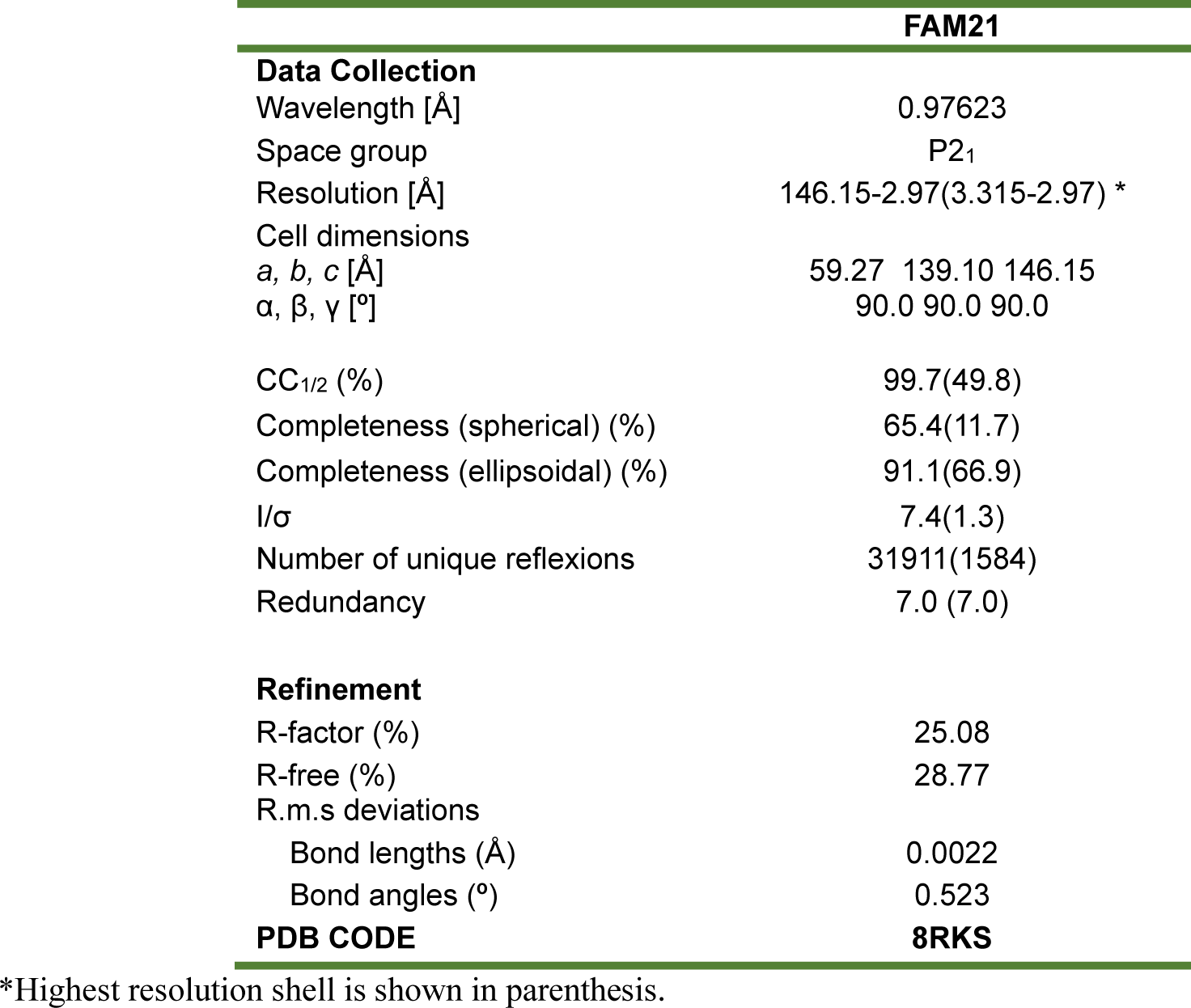
Summary of X-ray crystal data collection, phasing, and refinement statistics for the retromer complex bound to the R_21_ motif of Fam21.

### Structural comparison of R_21_ with other VPS29-effector complexes

Previous structural studies have uncovered the binding mode of other VPS29 effectors such TBC1D5 (22), VARP (5), VPS35L (15), and RidL (2; 44; 52). Superposition of the crystal structure of R_21_ with these effectors revealed a similar pattern for binding where a highly conserved proline, denoted as P_0_ position, represents the linchpin of the interaction (Figue 5A, B). The constrained geometry of this proline favors a sharper bend of the loop and introduces rigidity to the neighboring residues. In particular at position P_1_, there is a conserved leucine or isoleucine that, together with the proline at P_0_, form a conformationally restricted hydrophobic tip, optimizing the fit between R_21_ and the VPS29 pocket. Interestingly, the P-L amino acid tandem in R_21_ is unique among the 21 LFa repeats in Fam21 and agrees with the above results on its exclusive interaction with VPS29.

Given that VPS29, either alone or in complex with VPS35, has shown distinct affinities towards TBC1D5 (K_d_ ∼ 0.22 - 0.45 μM) (22; 44), VARP (K_d_ ∼ 2.7 – 5 μM) (5), VPS35L (K_d_ ∼ 1.87 μM) (15), RidL (K_d_ ∼ 0.15 - 0.5 μM) (2; 44; 52), and the R21 motif of Fam21(K_d_ ∼ 20 μM, this work), we examined the energy contribution at the residue level for each of the loops that contact the conserved hydrophobic pocket in VPS29 extending from amino acids P_-2_ to P_2_ (Figure 5C). With the exception of VARP, the remaining VPS29 effectors exhibited a similar pattern of energy contribution at each position, with P_-1_ to P_1_ identified as the hot-spot residues. Discrepancies with VARP may have arisen due to the methodological source of structures used for the analysis. While the energetic values for VARP originated from averaged NMR conformers that genuinely reflect the solution state, the remaining structures were derived from single crystallographic conformers, typically considered to adopt minimal energy landscapes within the lattice. In summary, this analysis indicates that the anchoring strength at the tip of each of the loops inserted into the hydrophobic pocket of VPS29 is very similar. The differences in affinity between distinct VPS29 effectors may therefore arise from the expansion of the interface area around the hydrophobic pocket or from the inclusion of additional intermolecular binding sites. As such, TBC1D5 contacts both VPS29 and the VPS35 subunits of retromer whereas the VPS35L subunit of retriever clamps VPS29 through the N- and C-terminal regions (15). On the other hand, RidL does not have additional intermolecular interactions with retromer but exhibits an extended buried surface next to the hairpin loop that further stabilizes the association with VPS29 (2; 44; 52). VARP, on the contrary, holds two Zn-fingernails with conformationally restrained scaffolds for binding VPS29 individually (18; 33; 5). In the case of Fam21, it employs the R_21_ motif exclusively for binding to VPS29 with low affinity through a relatively small buried interaction, while R_20_ and R_1-4_, on the other hand, associate with distinct sites on VPS35.

**FIGURE 5:**
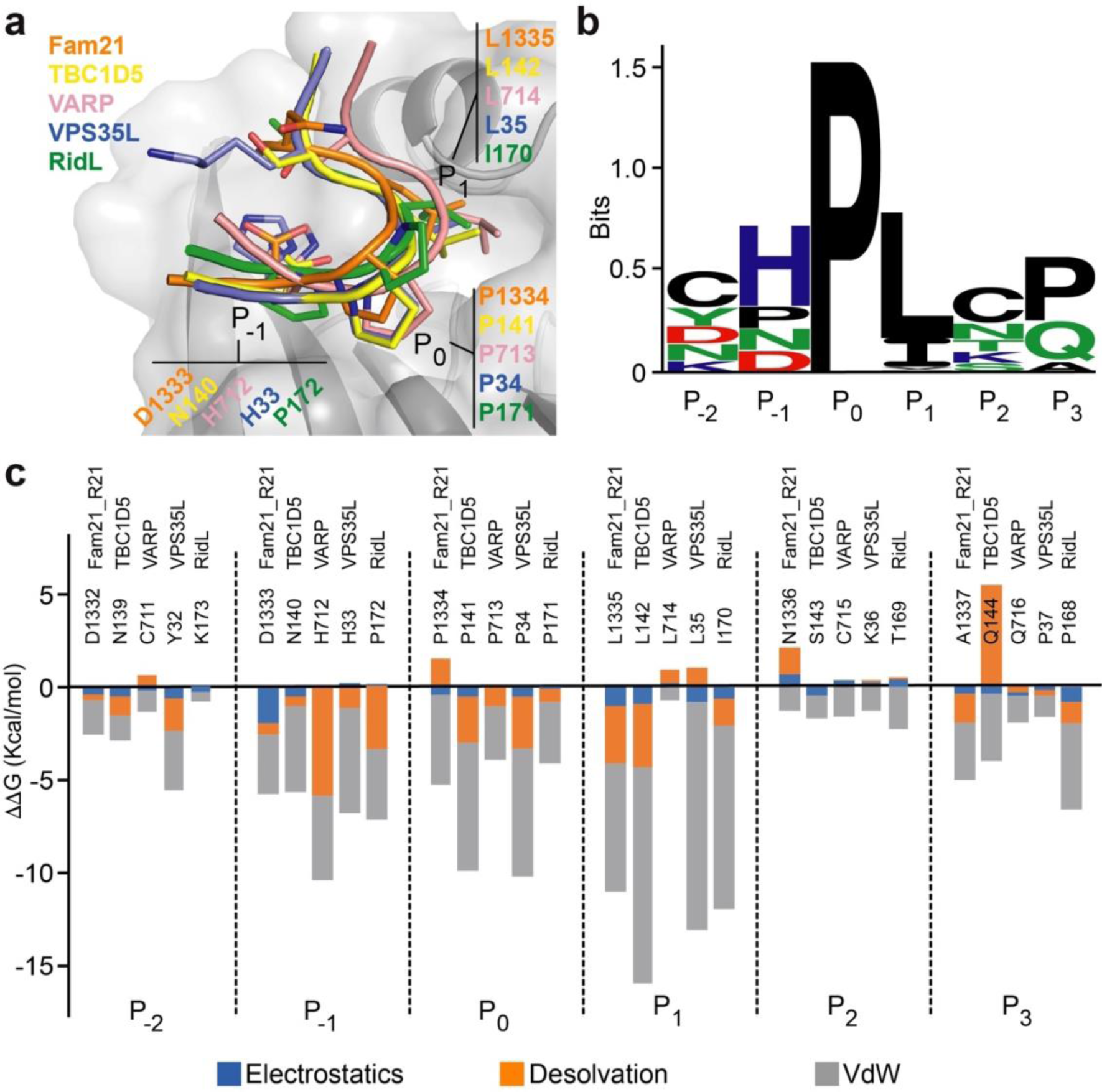
Structural comparison of the R_21_-VPS29 complex and other ligand complexes. (a) Superimposition of available structures from VPS29 ligands onto the VPS29 pocket. (b) WebLogo analysis of the superimposed structures. Note that despite RidL aligns in reverse direction, the tight P-L/I turn is conserved. (c) Comparative description of per-residue contributions from electrostatics (blue), van der Waals (vdW) (grey), and desolvation (orange) for residues inserted into the VPS29 pocket, extending from P-2 to P2.

## DISCUSSION

Since the identification of the 21 LFa repeats within the C-terminal unstructured tail of Fam21 (20), their multivalency and binding attributes have remained enigmatic. In the current study, we expanded the analysis of Fam21 and retromer interactions, identifying three distinct regions with diverse binding modes. While R20 and R21 bind to VPS35 and VPS29 subunits of retromer, respectively, as short linear motifs, the R_1-4_ segment, which contains four LFa repeats, requires the entire stretch of amino acids for binding to VPS35, indicating the presence of certain spatial constraints within this segment. In line with earlier findings (20), we observed that R_20_ and R_21_ contribute most significantly to the binding with retromer. However, despite previous indications of R_19_ playing a role in retromer binding (20), we couldn’t detect a measurable association when using the R_5-19_ fragment. In this regard, it is possible that R_19_ functions as a relatively weak binder in R_5-19_, while in the context of R_19-21_, it assumes a more prominent interaction with retromer, possibly due to inter-motif distances at which favorable binding occurs.

Previous studies indicated that Fam21 binds to the VPS35 subunit of the retromer complex (20; 16), and that this binding relies on the VPS35-VPS29 association(16). Here, we identified a direct interaction between Fam21 and VPS29 mediated by the R_21_ motif. This motif includes a distinctive P-L amino acid tandem, which is unique among the 21 LFa repeats. The crystal structure showed that the P-L tandem forms the tip of a tight turn that inserts into a conserved hydrophobic pocket on the surface of VPS29. This pocket is located opposite to the VPS29-VPS35C interface and is an important site for interaction with other endosomal effectors such as TBC1D5 (22), VARP(5), VPS35L(15), and RidL(2; 44; 52). Indeed, siRNA knockdown of TBC1D5 in HeLa cells leads to an increased interaction between retromer and the WASH complex (45). Interestingly, most of these effectors share a similar binding mode and per-residue energy contribution within the hydrophobic pocket, suggesting that the variations in affinity may arise from either the enlargement of the interface area surrounding the hydrophobic pocket and/or the incorporation of additional intermolecular binding sites. Retromer forms dimeric arch-like structures (Figure S5A,B) where the VPS26 subunit, in combination with distinct SNXs and cargo proteins, localizes at the base of the arch near the membrane surface (31; 25; 30). Meanwhile, VPS29 localizes away from the crowded membrane surface at the apex of the arch, reinforcing the idea of a central scaffold for the assembly of accessory proteins to establish distinct sorting stations on endosomes (1). In this context, the binding of R_21_ to VPS29, along with the interaction of both R_1-4_ and R_20_ with the C-terminal half of VPS35, occurs away from the membrane surface (Figure S5C), potentially facilitating the recruitment of the WASH complex to endosomes. Interestingly, the D620N mutation in VPS35, linked to PD, is localized at the apex where VPS35 homodimerizes. This mutation did not significantly reduce the direct binding towards R_1-4_ or R_20_ *in vitro*, suggesting that the perturbed association with Fam21 and the WASH complex (34; 54; 6) might arise from indirect effects or changes in the dynamic behavior of the retromer complex.

Our observations define discrete interactions between the unstructured tail of Fam21 and retromer. Importantly, the results align with the notion that the LFa motifs in FAM21, rather than promoting local clustering of retromer through tandem repetitions, might instead enable specific interactions with a number of factors using minor sequence modifications (8). We speculate that the consensus L-F-[D/E]3-10-L-F motif could display specificity through defined sequences and spatial distribution. As such, the spacing between the two L-F tandem repeats within each LFa motif, using variable stretches of negatively charged residues, could fine-tune their binding specificities. Meanwhile, the global positioning of the 21 LFa repeats could serve as an ’adaptor’ system where inter-motif distances might favor networks of interactions for the establishment of sorting subdomains. Future work should provide valuable information about the individual LFa binders and how such interactions evolve over time.

## MATERIALS AND METHODS

### Recombinant DNA procedures

DNA sequences encoding different fragments of FAM21 (R_1-19_, R_1-21_, R_1-4_, R_20- 21_, R_20_, R_21_, R_5-19_, R_1-2_, R_2-4_, R_3-4_) and retromer subunit VPS35C_204-780_ were cloned by Gibson assembly into pGST-Parallel2 vector (46) with a cleavable N-terminal glutathione S-transferase (GST) tag (19). Site-directed mutations in FAM21 R_20_, FAM21 R_21_ and VPS35 coding sequences, were introduced using the Phusion Site-Directed Mutagenesis Kit (Thermo Fisher) according to the manufacturer’s directions. All constructs were verified by DNA sequencing. For the expression of retromer complex (VPS29-VPS35- VPS26), the following plasmids were used: pMR101A-VPS29 (19), pGST-Parallel2- VPS35 (19) and pET28-Sumo3-VPS26 (31). In order to express the amino or carboxil terminal fragments of VPS35, pGST-Parallel2-VP35N (31) and pGST-Parallel2-VPS35C (19) vectors were used. For the expression of VPS29 mutants on Y165, Y163 and L152, the following vectors were used: pGST-Parallel2-VPS29, pGST-Parallel2-VPS29- Y165A, pGST-Parallel2-VPS29-Y163A and pGST-Parallel2-VPS29-L152E (44).

### Protein expression and purification

Proteins were expressed in E. coli BL21 (DE3) cells and grown in Luria–Bertani (LB) broth supplemented with the appropriate antibiotic at 37 °C. Protein expression was induced at an OD600 of 0.8 by adding 1 mM isopropyl-β-D-thiogalactopyranoside (IPTG). Cells were harvested after 16 h of growth at 18 °C. Purification steps were carried out at 4 °C, and the final protein concentration was determined using the theoretical extinction coefficient.

For the purification of VPS35 and FAM21 constructs, the cellular pellet was lysed using high-pressure homogenization (25 kpsi) in 50 mM Tris-HCl pH 7.5, 300 mM NaCl, 1 mM dithiothreitol (DTT) (buffer A), supplemented with 0.1 mM phenylmethylsulfonyl fluoride (PMSF) and 1 mM benzamidine. After clearing the bacterial lysates by centrifugation for 45 min at 50,000 × g, the soluble fraction was incubated in batch for 2 h with glutathione Sepharose 4B (GE Healthcare), previously equilibrated in 50 mM Tris-HCl pH 7.5, 150 mM NaCl, 1 mM DTT (buffer B). Protein was released from the resin by overnight cleavage of the N-terminal GST tag with TEV protease in buffer B. Subsequently, ion-exchange chromatography (HiTrap Q HP) was performed using a linear gradient of 20–1,000 mM NaCl, followed by size-exclusion chromatography on a Superdex 75 16/60 or Superdex 200 16/60 column based on the molecular weight of the protein, in buffer B. For the purification of VPS29 constructs, we followed the procedure described in Romano-Moreno et al., 2017 (44).

To purify the different protein complexes in all cases, cell pellets were mixed and purified together following the procedures described in Lucas et al., 2016 (31), for the retromer complex (VPS35-VPS29-VPS26) and the amino-terminal complex part (VPS35N-VPS26). For the purification of the carboxyl-terminal part (VPS35C-VPS29), the procedure described in Hierro et al., 2007 (19), was used. In the case of the extended carboxyl complex purification (VPS35_204-780_-VPS29), the procedure described in Hierro et al., 2007, was also employed.

The peptide FAM21-R_21_ (_1328_SNIFDDPLNAFGGQ_1341_) was synthesized by GenScript with a purity percentage greater than 98%.

### Protein crystallization, data collection and structure determination

The VPS35_476-780_-VPS29-FAM21 R_21_ complex was crystallized using hanging drop vapor diffusion method. Crystals were obtained by mixing the VPS35_476-780_-VPS29 complex at 143 µM with an excess of the FAM21 R_21_ peptide (_1328_SNIFDDPLNAFGGQ_1341_) at 1.5 mM. Native crystals appeared after 3 days at 18 °C, by mixing 1ul of protein sample with 1ul of the precipitant solution containing 20% (w/v) PEG 3350, 0.1 M NaCl and 0.1 M Tris-HCl pH 8.5. Crystals were cryoprotected by quick- soaking into mother liquor supplemented with 25% (v/v) ethylene glycol before being flash-frozen in liquid nitrogen.

Crystallographic native datasets were collected with the software MxCuBE at XALOC beamline in the ALBA synchrotron facility (Cerdanyola del Valles, Spain) using a Pilatus 6 M detector, and at Diamond Light Source (Oxfordshire, UK) with the software GDA. Diffraction images were indexed, integrated and scaled using XDS software (23). The VPS35_476-780_-VPS29-FAM21 R_21_ structure was solved by molecular replacement using as a template the complex VPS29-VPS35C (PDB 2R17) in PHASER (32).

The asymmetric unit contained four copies of VPS35_476-780_-VPS29, with one FAM21 R_21_ molecule bound to each VPS29. Peptide sequence was manually built in COOT (9) through iterative refinement with REFMAC5 (38). Data collection statistics for each dataset are shown in Table 1.

### Isothermal titration calorimetry

ITC measurements were carried out at 25 °C on a MicroCal PEAQ-ITC titration microcalorimeter (Malvern Panalytical). All proteins and peptides used in this work were dialyzed overnight at 4 °C against 300 mM NaCl, 0.5 mM TCEP, 25 mM HEPES pH 7.5 buffer. Before titration, samples were tempered at 25 °C and degassed for 5 min in a Thermo Vac. The titration sequence consisted of an initial 2µl injection to prevent artifacts (not used in data fitting), followed by 18 injections of 2 s and 2 µl with a spacing of 150 s between them. Heat of dilution used to correct the experimental data was performed under the same conditions. Results were fitted and integrated to a one-site model using the MicroCal PEAQ-ITC software (Malvern Panalytical). Final graphs were prepared using Origin ITC software (MicroCal). Values for the binding constant (Ka, Kd = 1/Ka), the molar binding stoichiometry, binding enthalpy, free energy and entropy of binding were obtained after data analysis. For ITC analysis of the interaction between different FAM21 motifs and retromer, 250 µM of FAM21 R_1-19_, R_1-21_, R_1-4_, R_20-21_, R_20_, R_21_, R_5-19_, R_1-2_, R_2-4_, R_3-4_ and the corresponding mutants, were titrated into 10 µM of retromer complex. For studying the region of retromer involved in the interaction with FAM21 regions R_1-4_, R_20_ and R_21_, 250 µM of each region was titrated into 10 µM of retromer, VPS35, VPS26-VPS35N, VPS29-VPS35C, VPS29-VPS35_ext_, or VPS35C, VPS29 and its different point mutants. The data are representative of a minimum of two replicate titrations for each assay.

### Computation of per-residue docking energy

We estimated the residue contribution to the binding energy using the pyDockEneRes Server (https://life.bsc.es/pid/pydockeneres) with default parameters. The input PDBs for VPS29-TBC1D5, VPS29-VPS35L, and VPS29-RidL crystal structures were 5GTU, 8ESE, and 5OSH, respectively. For the VPS29-VARP complex (6TL0), the energy contribution was calculated based on the average of ten NMR ensembles.

## SUPLEMENTARY MATERIAL DESCRIPTION

Supplementary material includes five figures and two tables

## Supporting information

Supplementary Material Romano-Moreno et al.

## ACKNOWLEDGMENTS

This study made use of the Diamond Light Source (Oxfordshire, UK) proposal MX20113, and ALBA synchrotron beamline BL13-XALOC. We express our gratitude to all beamline staff for their valuable assistance.

## DATA AVAILABILITY STATEMENT

Atomic coordinates and structure factors of the crystallographic complexes are available in the Protein Data Bank (PDB) with accession code 8RKS listed in Table 1.

## AUTHOR CONTRIBUTIONS

M.R.M. performed cloning, purification, crystallization, structure solution and ITC assays. E.N.A.S. performed purification, ITC assays and computational analysis of binding free energies. A.L.R. performed X-ray crystal structure solution. A.H. designed the research, analyzed the data and wrote the manuscript with the help of M.R.M. and E.N.A.S.

