## Supplementary Material Romano-Moreno et al. for "Retromer-mediated recruitment of the WASH complex involves discrete interactions between VPS35, VPS29 and Fam21"

**Affiliantions**

[1]Center for Cooperative Research in Biosciences (CIC bioGUNE), Bilbao, Spain

[2]Current address: i+Med S. Coop., Parque Tecnológico de Álava, Albert Einstein 15, nave 15, 01510 Vitoria-Gasteiz, Spain.

[3]IKERBASQUE, Basque Foundation for Science, Bilbao, Spain

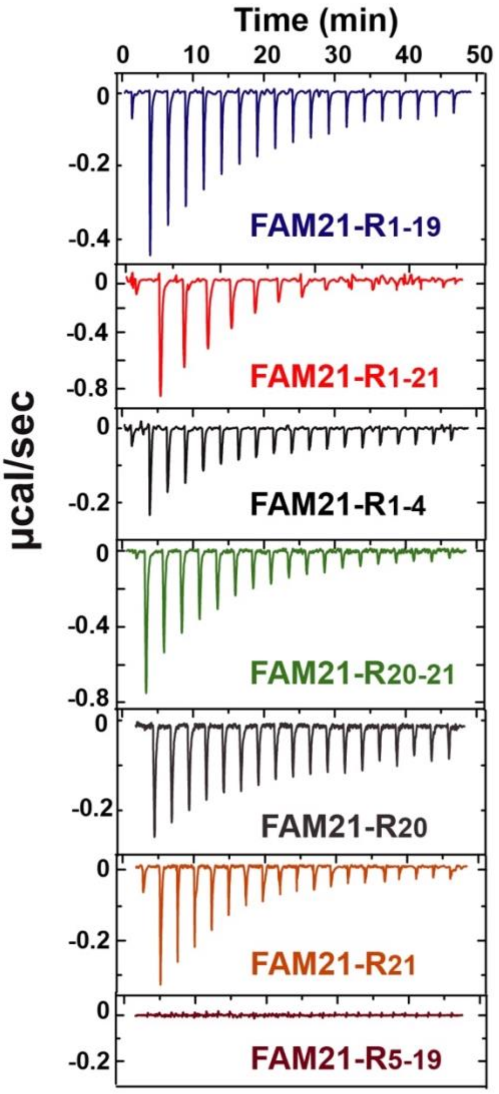


**SUPPLEMENTAL FIGURE 1:** Raw data from representative ITC measurements for titration of distinct segments of the Fam21 tail into retromer.

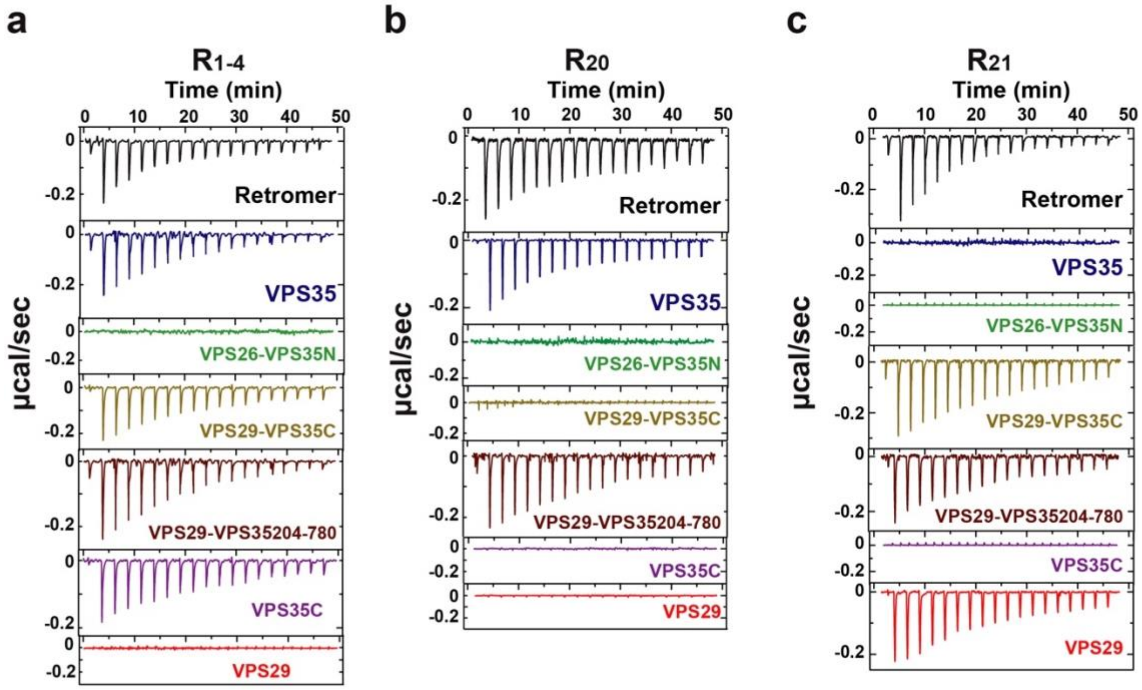


**SUPPLEMENTAL FIGURE 2:** Raw data from representative ITC measurements for titration of (a) $R_{1-4}$ fragment, (b) $R_{20}$ fragment, and (c) $R_{21}$ fragment of Fam21 to individual retromer subunits or truncated subcomplexes.

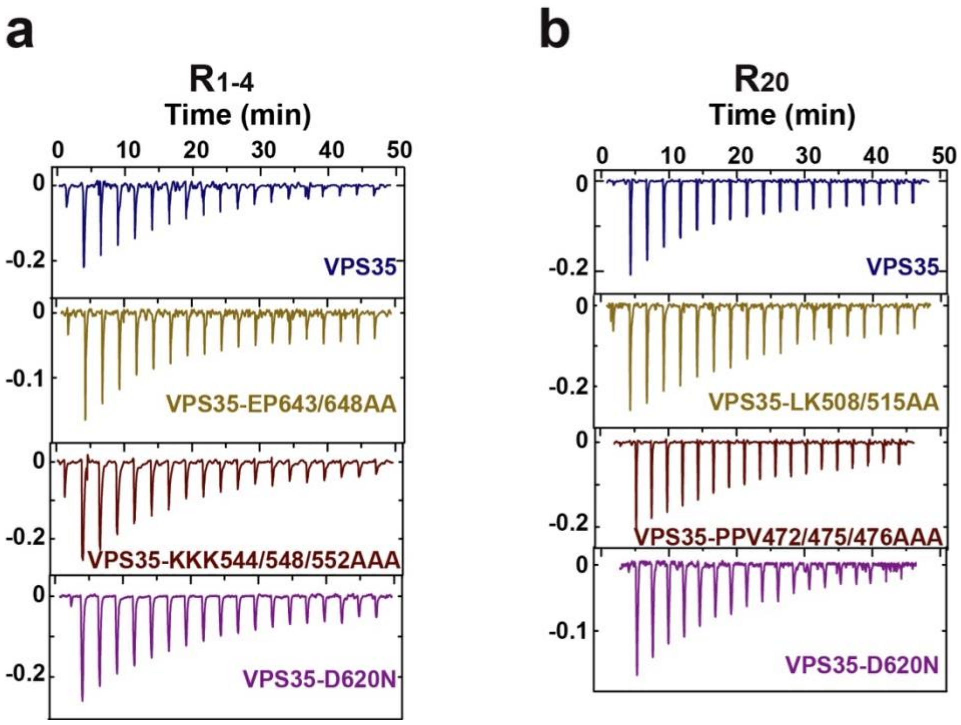


**SUPPLEMENTAL FIGURE 3:** Raw data from representative ITC measurements for titration of (a) $R_{1\text{-}4}$ fragment, and (b) $R_{20}$ fragment of Fam21 to wildtype VPS35 subunit and selected mutants.

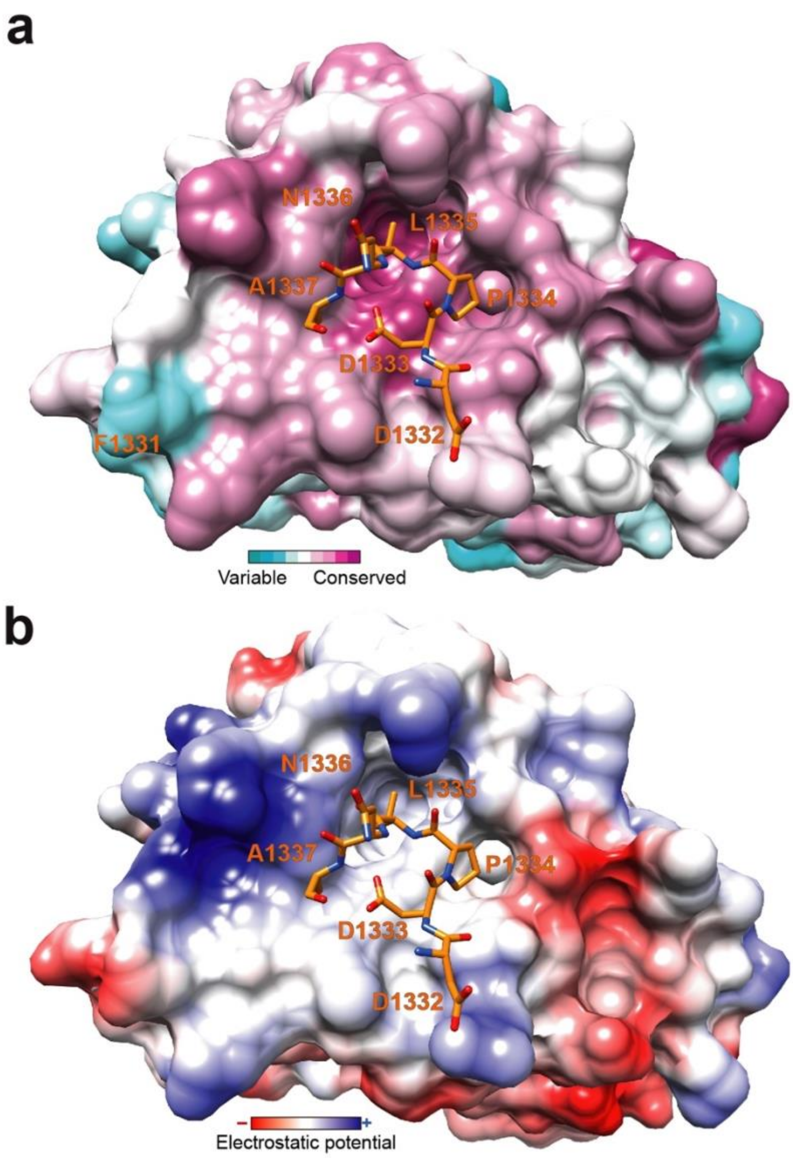


**SUPPLEMENTAL FIGURE 4:** (a) Conserved accessible surface of VPS29 colored according to xProtCAS(26). The color-coding legend is provided below, and the R21 motif is depicted as a stick model in orange. (b) Electrostatic surface potential (−10 to +10 kcal/mol·e in red to blue) mapped on the surface of VPS29 structure, in the same orientation as in (a).

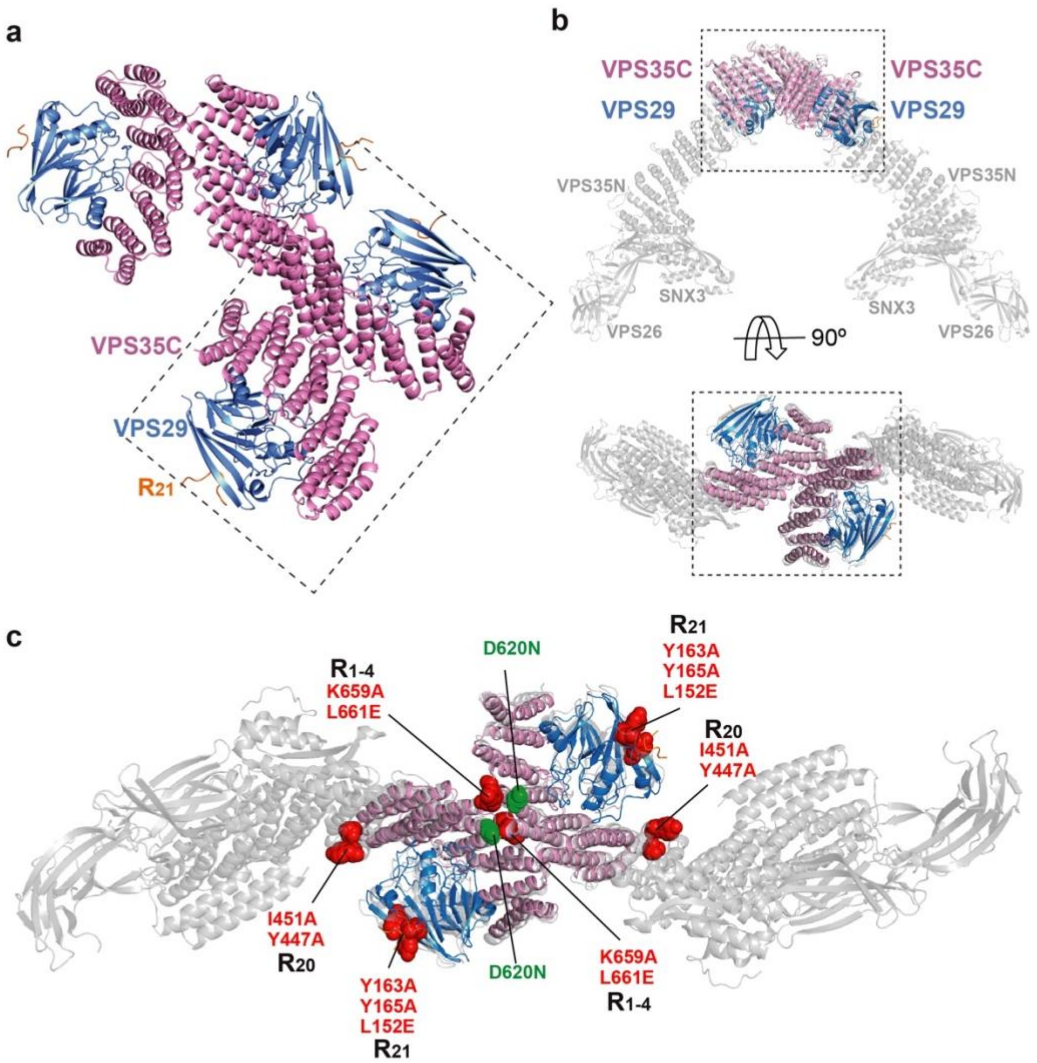


**SUPPLEMENTAL FIGURE 5:** (a) Packing of four VPS35C-VPS29-R21 complexes that form an asymmetric unit. Dashed box highlights a crystal contact between two VPS35C molecules mimicking the dimeric interface observed in the *membrane*-assembled retromer coat (32). (b) superposition of the membrane assembled retromer coat with one of the two dimeric units of VPS35C-VPS29-R21 present in the crystallographic asymmetric unit. (c) Mutations performed in the current work, highlighted in red within the membrane-assembled retromer (view from the bottom of the arch), that precluded the interaction with $R_{1-4}$, $R_{20}$, and $R_{21}$. In green, the Parkinson's disease-linked mutation (D620N).

**SUPPLEMENTAL TABLE 1:** Summary of ITC thermodynamic parameters.

| | Kd (μM) | n | ΔH (kcal/mol) | -TΔS (kcal/mol) | ΔG (kcal/mol) |
|---|---|---|---|---|---|
| **FAM21-R1-19 + [VPS26-VPS29-VPS35]** | 20 ±3 | 0.7 ±0.2 | -13.4 ±4 | 7.6 ±2.4 | -5,8 ±1.6 |
| **FAM21-R1-21 + [VPS26-VPS29-VPS35]** | 4 ±1 | 0.4 ±0.1 | -20.8 ±3 | 13.5 ±1.3 | -7.3 ±1.6 |
| **FAM21-R1-4 + [VPS26-VPS29-VPS35]** | 21 ±3 | 1 ±0.3 | -17.4 ±5 | 11.1 ±4.2 | -6.3 ±0.6 |
| **FAM21-R20-21 + [VPS26-VPS29-VPS35]** | 9 ±1 | 0.5 ±0.02 | -17.5 ±1.5 | 11.4 ±0.2 | -6.4 ±1.3 |
| **FAM21-R20 + [VPS26-VPS29-VPS35]** | 20 ±1 | 1.1 +±0.2 | -16.8 ±2 | 9.3 ±0.5 | -7.5 ±1.6 |
| **FAM21-R21 + [VPS26-VPS29-VPS35]** | 19 ±3 | 1 ±0.1 | -17.8 ±1.5 | 10.6 ±0.6 | -7.2 ±0.8 |
| **FAM21-R21L1335E + [VPS26-VPS29-VPS35]** | N.B. | | | | |
| **FAM21-R20IF1290/1291AA + [VPS26-VPS29-VPS35]** | N.B. | | | | |
| **FAM21-R20IF1298/1299AA + [VPS26-VPS29-VPS35]** | N.B. | | | | |
| **FAM21-R1-4 + VPS35** | 20 ±2 | 1 ±0.1 | -11.2 ±5 | 4.8 ±1.2 | -6,4 ±3.6 |
| **FAM21-R1-4 + VPS26-VPS35N** | N.B. | | | | |
| **FAM21-R1-4 + VPS29-VPS35C** | 23 ±4 | 0.7 ±0.2 | -7.4 ±0.6 | 1.1 ±0.2 | -6,3 ±0.3 |
| **FAM21-R1-4 + VPS29-VPS35204-780** | 21 ±1 | 1.2 ±0.1 | -8,3 ±1 | 2.8 ±0.6 | -5.4 ±1.6 |
| **FAM21-R1-4 + VPS35C** | 47 ±6 | 1 ±0.1 | -6.5 ±0.5 | -1.4 ±0.4 | -5.9 ±0.1 |
| **FAM21-R1-4 + VPS29** | N.B. | | | | |
| **FAM21-R20 + VPS35** | 20 ±1 | 1 ±0.1 | -5.2 ±0.5 | 1.9 ±0.3 | -3.2 ±0.8 |
| **FAM21-R20 + VPS26-VPS35N** | N.B. | | | | |
| **FAM21-R20 + VPS29-VPS35C** | N.B. | | | | |
| **FAM21-R20 + VPS29-VPS35204-780** | 21 ±1 | 1.1 ±0.1 | -4.0 ±1 | 1.2 ±0.3 | -2.8 ±0.6 |
| **FAM21-R20 + VPS35C** | N.B. | | | | |
| **FAM21-R20 + VPS29** | N.B. | | | | |
| **FAM21-R21 + VPS35** | N.B. | | | | |
| **FAM21-R21 + VPS26-VPS35N** | N.B. | | | | |
| **FAM21-R21 + VPS29-VPS35C** | 19 ±2 | 1.2 ±0.1 | -7.2 ±0.9 | 1.6 ±0.1 | -5,6 ±0.7 |
| **FAM21-R21 + VPS29-VPS35204-780** | 19 ±3 | 1.1 ±01 | -4.5 ±1 | 1.0 ±0.3 | -3.4 ±0.6 |
| **FAM21-R21 + VPS35C** | N.B. | | | | |
| **FAM21-R21 + VPS29** | 21 ±3 | 1 ±0.1 | -6.1 ±0.5 | -0,3 ±0.2 | -6.4 ±0.3 |

**SUPPLEMENTAL TABLE 2:** Key resources

| REAGENT or RESOURCE | SOURCE | IDENTIFIER |
|---|---|---|
| **Chemicals, Peptides, and Recombinant Proteins** | | |
| Peptide FAM21 $R_{21}$ $_{1328}$SNIFDDPLNAFGGQ$_{1341}$ | GenScript | N/A |
| **Critical Commercial Assays** | | |
| Glutathione Sepharose 4B | GE Healthcare | Cat# 17-0756-05 |
| Ni-NTA Agarose | Qiagen | Cat# 30230 |
| HiTrap Q HP 5ml column | GE Healthcare | Cat# 17-1154-01 |
| HiLoad 16/60 Superdex 75 column | GE Healthcare | Cat# 17-1068-01 |
| HiLoad 16/60 Superdex 200 column | GE Healthcare | Cat# 17-1069-01 |
| **Deposited Data** | | |
| VPS35C-VPS29-FAM21-$R_{21}$ | This study | PDB: 8RKS |
| **Experimental Models: Organisms/Strains** | | |
| *Escherichia coli* BL21(DE3) | Invitrogen | Cat# C600003 |
| *Escherichia coli* DH5α | Invitrogen | Cat# 18258012 |
| **Recombinant DNA** | | |
| pGST-Parallel2 | (49) | N/A |
| pET28-Sumo3-VPS26 | (33) | N/A |
| pMR101A-VPS29 | (19) | N/A |
| pGST-Parallel2-VPS29 | (47) | N/A |
| pGST-Parallel2-VPS29-Y165A | (47) | N/A |
| pGST-Parallel2-VPS29-Y163A | (47) | N/A |
| pGST-Parallel2-VPS29-L152E | (47) | N/A |
| pGST-Parallel2-VPS35 | (19) | N/A |
| pGST-Parallel2-VPS35$_{(14-470)}$ (VPS35N) | (33) | N/A |
| pGST-Parallel2-VPS35$_{(476-780)}$ (VPS35C) | (19) | N/A |
| pGST-Parallel2-VPS35C$_{(204-780)}$ | This study | N/A |
| pGST-Parallel2-VPS35-KL659/661AE | This study | N/A |
| pGST-Parallel2-VPS35-EP643/648AA | This study | N/A |
| pGST-Parallel2-VPS35-KKK544/548/552AAA | This study | N/A |
| pGST-Parallel2-VPS35-D620N | This study | N/A |
| pGST-Parallel2-VPS35-YI447/451AA | This study | N/A |
| pGST-Parallel2-VPS35-LK508/515AA | This study | N/A |
| pGST-Parallel2-VPS35-PPV472/475/476AAA | This study | N/A |
| pGST-Parallel2-FAM21-$R_1$-$R_{21}$ | This study | N/A |

| | | |
|---|---|---|
| pGST-Parallel2-FAM21-$R_1$-$R_{19}$ | This study | N/A |
| pGST-Parallel2-FAM21-$R_5$-$R_{19}$ | This study | N/A |
| pGST-Parallel2-FAM21-$R_1$-$R_4$ | This study | N/A |
| pGST-Parallel2-FAM21-$R_1$-$R_2$ | This study | N/A |
| pGST-Parallel2-FAM21-$R_2$-$R_4$ | This study | N/A |
| pGST-Parallel2-FAM21-$R_3$-$R_4$ | This study | N/A |
| pGST-Parallel2-FAM21-$R_{20}$-$R_{21}$ | This study | N/A |
| pGST-Parallel2-FAM21-$R_{20}$ | This study | N/A |
| pGST-Parallel2-FAM21-$R_{20}$-IF1290/1291AA | This study | N/A |
| pGST-Parallel2-FAM21-$R_{20}$- IF1298/1299AA | This study | N/A |
| pGST-Parallel2-FAM21-$R_{21}$ | This study | N/A |
| pGST-Parallel2-FAM21-$R_{21}$-L1335E | This study | N/A |
| **Software and Algorithms** | | |
| XDS | (23) | http://xds.mpimf-heidelberg.mpg.de |
| CCP4 | (55) | http://www.ccp4.ac.uk |
| COOT | (9) | http://www2.mrc-lmb.cam.ac.uk/personal/pemsley/coot/ |
| UCSF CHIMERA | (44) | https://www.cgl.ucsf.edu/chimera/uc |
| PYMOL | Molecular Graphics System, Version 1.8 Schrödinger, LLC | https://www.pymol.org/ |
| MicroCal PEAQ-ITC software | Malvern Panalytical | https://www.malvernpanalytical.com/es/support/product-support/software/microcal-peaq-itc-analysis-software-v141 |
| Origin ITC software | MicroCal | https://www.malvernpanalytical.com/en/support/product-support/software/microcal-itc-origin-add-on-dissociation-model-update-v1-00 |
| pyDockEneRes Server | Barcelona Supercomputing Center | https://life.bsc.es/pid/pydockeneres |
| LigPlot+ | (30) | https://www.ebi.ac.uk/thornton-srv/software/LigPlus/ |
